# Assessing the utility of MAGNETO to control neuronal excitability in the somatosensory cortex

**DOI:** 10.1101/762559

**Authors:** Koen Kole, Yiping Zhang, Eric J. R. Jansen, Terence Brouns, Ate Bijlsma, Niccolo Calcini, Xuan Yan, Angelica da Silva Lantyer, Tansu Celikel

**Affiliations:** Department of Neurophysiology, Donders Institute for Brain, Cognition and Behaviour, Radboud University, the Netherlands

## Abstract

Magnetic neuromodulation has outstanding promise for the development of novel neural interfaces without direct physical intervention with the brain. Here we tested the utility of Magneto in the adult somatosensory cortex by performing whole-cell intracellular recordings in vitro and extracellular recordings in freely moving mice. Results show that magnetic stimulation does not alter subthreshold membrane excitability or contribute to the generation of action potentials in virally transduced neurons expressing Magneto.

---

Recently introduced Magneto (Wheeler et al., 2016) might provide the highly sought after neuromagnetic actuation in a cell-targeted manner. Some of the excitement about Magneto originates from its design which is comprised of a calcium-permeable non-selective cation channel (Transient receptor potential cation channel subfamily V member 4, TRPV4) fused to the paramagnetic protein ferritin (Wheeler et al., 2016). This single-construct approach provides a simplified mean for magnetic intervention with neuronal activity. Here, we used lentiviral delivery of Magneto linked to mCherry (Magneto2.0-P2A-mCherry), expressed under the control of ubiquitin promoter for >2 weeks **(Fig.1a)** before observing and interfering with neural activity (see Methods online), and after confirming successful cleavage of Magneto from mCherry **(Suppl.Fig.2a-3)** and the subcellular analysis of the expressed protein localization **(Suppl.Fig.2b)** in a neuronal cell line. Chronic extracellular recordings in freely moving mice (Allen et al., 2003; Celikel et al., 2004; Clem et al., 2008) with 15 tetrodes enabled high-density sampling of neural activity in the vicinity of transduced cells, and yielded well-isolated **(Suppl.Fig.4)**, stable units **(Fig.1b)**. Comparison of firing rates within cells across magnetic stimulus conditions (off vs on) showed that magnetic stimulation does not alter the rate of action potentials (APs; **Fig.1c**); neither does it modulate the inter-spike interval within cells, nor spike-timing across single units recorded from the same tetrode **(Suppl.Fig.5,6)**. The lack of spiking was not because neurons could not respond to synaptic depolarization; deflection of magnetized whiskers (with nano iron particles, see Methods online) using an electromagnet (Clem et al., 2008), induced stimulus-coupled spiking **(Fig.1d)**. Considering that direct neuromagnetic stimulation failed to trigger any change in low-frequency local-field potential oscillations **(Fig.1e,f)**, we argue that the utility of Magneto to control neural activity in vivo is not warranted.

**Figure 1.**
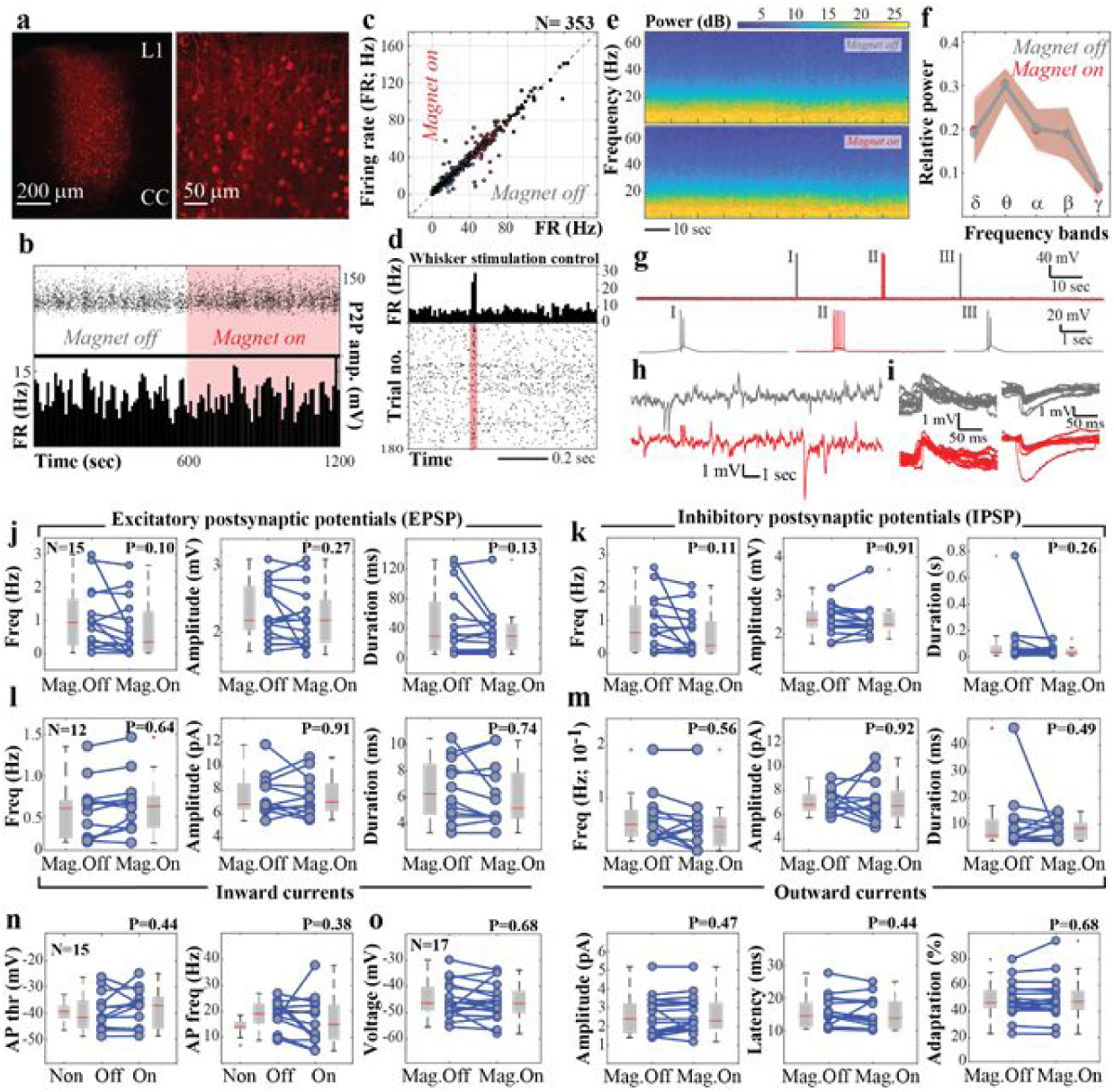
Electrical characterization of the consequences of magnetic neural stimulation. Neurons in the primary somatosensory cortex (barrel field) were transduced using a Lentiviral vector encoding Magneto-p2A-mCherry for at least two weeks before extracellular recordings with chronically implanted distributed microelectrode arrays, and whole-cell current and voltage clamp experiments were performed. Magnetic stimulation was provided with permanent magnets with an estimated magnetic field intensity of >50 mT in the vicinity of the cells recorded (see Supplemental Figure 1 for magnetic field strength measurements; see Methods Online for magnet placement). (a) Example of viral transduction throughout a cortical column of interest. Right: A higher magnification view from the cortical layers 2/3. (b) Rate of action potentials (bottom) and unit stability (top), as quantified by peak-to-peak (P2P) amplitude, during a 20 min long recording session for a single unit recording before and during magnetic stimulation in freely behaving mice. (c) Firing rate of 353 single units recorded across five sessions in two mice. The two axes denote the firing rates for each neuron with or without magnetic stimulation. Magnetic stimulation did not alter the probability of action potential generation (P=0.64; paired t-test). See Supplemental Figure 7 for the standard deviation across stimulus conditions for each unit. (d) Raster plot and peristimulus time histogram of action potentials evoked by whisker stimulation delivered using an electromagnet after coating with a whisker with iron nanoparticles. (e) Power spectrogram of the local field potential (LFP) with or without magnetic stimulation (N= 24/condition). (f) Relative LFP power of across δ (1-4 Hz), θ (4-8 Hz), α (8-12 Hz), β (13-30 Hz), γ (30-70 Hz) bands. Magnetic stimulation did not change the LFP frequency or power (N=24/condition; P>0.99 for interaction between magnetic stimulation and LFP power across frequencies, two-way ANOVA). (g-i) Sample traces showing spiking (g), subthreshold activity (h) isolated excitatory and inhibitory postsynaptic potentials (i) and with (red) and without magnetic stimulation (grey) recorded in current-clamp configuration. Note that initiation of (bursts of) action potentials and subthreshold potentials lack temporal correlations with initiation and termination of magnetic stimulation, suggestive of spontaneous events. (j-k) Frequency, amplitude, and duration of the excitatory (j) or inhibitory (k) postsynaptic potentials are not modulated by magnetic stimulation (paired t-test, P values are on the figurines). For action potential statistics refer to the main text. (l-m) Spontaneous inward and outward currents were unaffected upon magnetic stimulation (paired t-test). (n) Analysis of action potential statistics in vitro. All statistical comparisons were performed using paired t-test within cells across magnetic stimulation conditions, i.e. “Off” (Magnet Off) vs “On” (Magnet On) in Magneto-expressing cells. Data from control cells did not express Magneto are labeled as “Non”. (o) Voltage-gated conductances as visualized using triangular voltage sweeps (see Supplemental Methods online for details). Rate of adaptation is quantified as the fractional change in the half-width of inward rectification (see Methods for details). In all box-plots center lines represent the distribution median and box limits are upper and lower quartiles; whiskers are 1.5x of the interquartile range and outliers are shown as crosses. These results argue that Magneto does not alter neuronal excitability or control action potential generation *in vivo* or *in vitro*.

Extracellular recordings with multi-electrode arrays enable high-throughput observation of neural activity without selectively targeting virally transduced neurons. If Magneto-mediated neuroactuation or viral transduction were limited in efficacy, the spatiotemporally correlated membrane depolarization across local populations triggered by magnetic stimulation would be insufficient to induce spiking in a large population of neurons, potentially explaining the negative results described above. Therefore we next performed intracellular recordings in visualized neurons that express the reporter fluorescence protein, mCherry **(Fig.1g-o)**. Whole-cell single unit recordings showed that magnetic stimulation does not change the probability of AP generation (Magnet-off: 0.7±1.4, Magnet-on: 0.7±1.8 spikes (mean±std); N=15 cells; P>0.8, paired t-test). Current-clamp and voltage-clamp recordings showed that magnetic stimulation is also ineffective in modulating the frequency, amplitude or duration of the subthreshold postsynaptic potentials **(Fig.1j,k)** and inward/outward currents **(Fig.1l,m)**, suggesting that magnetic stimulation alone is not sufficient to control neural activity in Magneto expressing neurons. Analysis of the action potential threshold and frequency of action potentials concluded that magnetic stimulation does not change the basic statistics of spiking in vitro **(Fig.1n)**. To address whether voltage-gated conductances might be independently modulated during magnetic stimulation, we performed current clamp recordings with triangular (sawtooth) voltage sweeps. The results showed that the membrane voltage at which the channels open, the amplitude and latency of inward rectification, and the rate of amplitude adaptation were comparable across magnetic stimulation conditions (magnet-off vs magnet-on) **(Fig.1o)**. These results argue that magnetic stimulation of neurons that express Magneto does not result in sub- or suprathreshold modulation of neural activity.

Neuromodulation via extracranial stimulation has outstanding promise for sensory and motor prosthetics applications. Considering the negligible expression of native TRPV channels in the barrel cortex ((Kole et al., 2017a, 2017b) for data visualization visit http://barrelomics.science.ru.nl), heterologous expression of TRPV4-ferritin might be considered as a suitable tool for controlled reproducible activation of somatosensory neurons. However, we failed to trigger action potentials or subthreshold depolarization by magnetic stimulation in vitro and in vivo. These findings replicate others’ observations made in Cerebellum (Xu et al., 2018), hippocampus, entorhinal cortex and barrel cortex (Wang et al., 2019), and in different cell types (N2A, current study, HEK293 (Wang et al., 2019)) using multiple expression systems (including transfection, lentivirus, Sindbis, AAV, FSV); some of these experiments were performed in the same neuronal classes, using an identical viral expression system and stimulation magnet as described in the original paper (Wheeler et al., 2016). In support of these findings, immunohistochemical observations **(Suppl.Fig.2b)** and biochemical measurements (Wang et al., 2019) show that Magneto2.0, upon expression, primarily remains in reticular structures in the cytoplasm and is not efficiently transported to the plasma membrane, although TRPV4 (i.e. Magneto2.0 primogenitor) is inserted into the membrane as expected **(Suppl.Fig.2b)**. A critical re-evaluation of the Magneto2.0 design might be necessary.

## Materials and Methods

All experimental procedures have been performed in accordance with the European Directive 2010/63/EU, guidelines of the Federation of European Laboratory Animal Science Associations, and the NIH Guide for the Care and Use of Laboratory Animals. Experiments were approved by the Animal Ethical Committee of the Radboud University Nijmegen, The Netherlands (permit numbers DEC-2013-172-001 and DEC-2014-275-001).

### Animals

Adult transgenic mice, B6;129P2-Pvalbtm1(cr)Arbr/J (N=6) or Ssttm2.1(cre)Zjh/J (N=1), were obtained from local breeding colonies, and maintained under *ad libitum* access to food and water, and 12/12h light/dark cycle. Animals were housed together with their littermates until the day of viral injection, after which they were housed individually to reduce the risk of postoperative injury. Animals that received drive implantation for chronic electrophysiological recordings were placed in larger cages with higher ceiling to reduce the risk of mechanical damage. In total 7 animals (3-7 months old) were used for the experiments described herein. Except 2 B6;129P2-Pvalbtm1(cr)Arbr/J mice used for chronic recordings, all mice were females.

### Lentiviral vector preparation

For in vivo gene delivery, pcDNA3.0-Magneto2.0-p2A-mCherry (Addgene #74308) was subcloned into lentiviral vector, pFUGW-V4trunc-fer-traffick-p2A-mCherry. Lentiviruses were produced as described before (Celikel et al., 2007) with modifications. In brief, HEK293T/17 (ATCC, CRL-11268) cells were seeded onto 8 × 10 cm dishes and per dish transfected with a total of 12 µg endotoxin-free plasmid DNA containing the helper plasmids pPL1 (3.4 µg), pPL2 (1.7 µg), pPL-VSVg (2.6 µg) and pFUGW-V4trunc-fer-traffick-p2A-mCherry (4.2 µg) using jetPRIME Transfection Reagent (Polyplus Transfection, #114-15) according to the manufacturer’s instructions. After 36 hours, culture medium (DMEM with 10% Fetal Calf Serum, 1mM Na-Pyruvate, 100 units/ml penicillin and 100 µg/ml streptomycin) was replaced with fresh medium containing 4 mM valproic acid (VPA). The next day, 24 hours later, the medium was removed from culture dishes, supplied with 20 mM Hepes and filtered using a prewashed 45 µm filter and fresh medium (containing VPA) was added to the dishes. Filtered medium was slowly transferred onto a cushion of 20% sucrose/Hank’s Balanced Salt Solution (HBSS) and ultracentrifuged for 5 hours at 12.500 rpm in a Sorvall Discovery 100 ultracentrifuge using an AH 629 rotor. After centrifugation, medium and sucrose/HBSS were discarded and the pellet was supplied with 50 µl HBSS and placed at 4°C overnight. This process was repeated with fresh medium 24 hours later, after which the second pellet was stored overnight at 4°C. The two pellets were then resuspended and combined to constitute a total of 200 µl suspension in HBSS, which was aliquoted and stored at -80°C until further use.

The genomic titers were determined using qPCR. Amplicons were generated using primers against WPRE (FW: 5’-ggcactgacaattccgtggt-3’, RV: 5’-agggacgtagcagaaggacg-3’; Sigma-Aldrich). To control for unpackaged plasmid DNA, viral suspensions were treated with DNAse I (Invitrogen, # 18068015). Each sample contained 5.5 µl milli-Q, 12.5 µl SensiFast SYBR-Green Mastermix (Bioline, #BIO-98005), 5 µM FW primer, 5 µM RV primer and 1 µl sample. An initial 2 minutes at 95 °C were followed by 40 cycles of melting (@95°C for 5 sec), annealing (@60°C for 10 sec), extension (@72°C for 20 sec). After cycling was complete, a melting protocol was performed; measuring fluorescence intensity from 60 to 95°C with a step size of 0.5°C to control for amplicon specificity. To determine the physical titer, a standard curve was generated based on the plasmid DNA. The calculated titer was 5.6 × 10^8^ particles/ml.

The nucleotide sequence of the Magneto 2.0-p2A-mCherry open reading frame in the lentiviral vector was verified by Sanger sequencing (Lightrun, GATC Biotech) using the following primers (Sigma-Aldrich):

Magneto-2FW (5’-caaggcacttctgaacttaagc-3’) Magneto-3FW (5’-ctggtttacaacagcaagatc-3’), Magneto-4FW (5’-ctggacctcttcaagctcac-3’) Magneto-5FW (5’-acttcctggagactcacttc-3’), Magneto-6FW (5’-tctttgacaagcacaccctg-3’) Magneto-7FW (5’-tcctccgagcggatgtac-3’), Magneto-1RV (5’-tagccaccctcatccttg-3’) Magneto-2RV (5’-ggagctccacgtaatgc-3’).

The sequence was compared against the pcDNA3.0-Magneto2.0-p2A-mCherry (Addgene #74308) using ContigExpress and AlignX (Vector NTI Advance 10, Invitrogen) which had 100% sequence similarity (see Supplemental Table 1).

### Viral gene delivery *in vivo*

Lentiviral particles were pressure injected as described before (Celikel et al., 2007; Freudenberg et al., 2013a, 2013b). For *in vitro* slice experiments (N=5 mice) in vivo viral injections were performed under isoflurane anaesthesia. Body temperature was monitored and maintained at 37±0.5 °C. A glass capillary with a ∼20 µm tip containing lentiviral particles was used to deliver the virus to the barrel subfield of the primary somatosensory cortex bilaterally (Anteroposterior: -1.5 mm from Bregma, and Mediolateral 3.0 mm from midline) after skin incision and retraction. Injections were initiated at a depth of ∼500 µm while slowly retracting the capillary until a depth of ∼200 µm was reached. A total of ∼300 nl was injected over 15 minutes, after which pneumatic pressure was removed and the capillary was left for an additional 2 minutes to allow viral particles to spread before full retraction of the injection needle. After the injections were completed, Carprofen (8-10 mg/kg) was administered subcutaneously. The skin was sutured and the animal was returned to its home cage after recovery. Animals were typically awake and mobile within 20 – 30 minutes.

For *in vivo* chronic recording experiments (N=2 mice), viral injections were performed through a polyimide tube (OD=105µm, ID=40µm) that was positioned at the center of 15 tetrodes carried by a “FlexDrive”, secured permanently to the skull. This ensured the spatial alignment of the virally transduced neurons and recording electrodes. Each tetrode was placed in a polyimide tube and connected a fine screw that allowed axial positioning of the tetrodes in the barrel cortex. The drive preparation was described as before(Voigts et al., 2013) with the exception of the inclusion of aforementioned access port for the viral injections. Viral injection was performed under isoflurane anesthesia as ∼300 nl viral vectors delivered in ∼15 minutes ∼200 *µ*m below the cortical surface.

### Chronic extracellular recordings

FlexDrive implantation was performed under isoflurane anesthesia while body temperature was maintained at 37±0.5 °C. The surgery started after subcutaneous injection of Cefazolin (20mg/kg), dexamethasone (2mg/kg), Carprofen (5mg/kg) and saline (15ml/kg) and intramuscular injection of Buprenorphine (0.03mg/kg). After skin incision, a window was prepared above the barrel cortex as described before (Allen et al., 2003; Celikel et al., 2004). In short, skull in a 4 mm^2^ area (0 to -2 mm from Bregma and 2 to 4 mm from midline) was thinned while intermittently cooling the skull using saline drops. Thinned skull was removed using a fine pair of forceps after the space between the skull and the pia was buffered with saline. The dura mater was incised using a 29G insulin syringe needle. The angle of the FlexDrive was adjusted to ensure perpendicular penetration of the electrodes into the cortex before the FlexDrive was fixed on the skull using dental cement (Super-Bond C&B). The animal was returned to a warm cage for recovery and returned to home cage where HydroGel and wet food were provided for ease of water and food consumption. Animals were awake within 30 minutes and regained motor activity shortly after.

One week after the surgery the electrodes were gradually inserted into the brain, moving the electrodes ∼50-200 *µ*m in every session (the speed is determined by the predicted distance to the brain and electrical recording characteristics in a given location). Data acquisition was performed using Open Ephys interface (Siegle et al., 2017) after the signals was filtered (0.1-6000 Hz) and digitized (@30 kHz/channel) using a 64 channel amplifier (Intan Technologies; RHD2164). The data was processed offline (see below). Magnetic stimulation during chronic recordings were provided either by a permanent magnet block, or a custom electromagnet described previously (Clem et al., 2008) after animals were habituated to the behavioral chamber for 15 minutes. The permanent magnet block consisted of three blocks (20×10×2 mm) and 14 ring (8 mm in diameter) neodymium magnets. It was manually placed within the 7-9 mm of the skull. Animals were not aware of the upcoming magnetic stimulation, as the experimental chamber was surrounded with translucent dark grey plexiglas walls (thickness = 3mm). The magnetic stimulation intensity at the recording site was calculated to be >50 mT (see Supplemental Figure 1 for magnetic intensity measurements). The results were comparable across magnetic stimulation conditions, and thus combined and presented together. Whisker deflections were delivered using the electromagnet after select individual whiskers in the C-row were coated with iron nanoparticles.

### Acute intracellular recordings

5 – 8 weeks post injection, acute brain slices were prepared as described before (Allen et al., 2003; Celikel et al., 2004; Clem et al., 2008) with modifications for the adult brain. In brief, ice cold slicing medium (Choline-Chloride 108 mM; KCl 3 mM; NaHCO3 26 mM; 6 MgSO4; NaHPO4 1.25 mM; D-glucose 25 mM; Na-pyruvate 3 mM; CaCl2 2 mM) was carbogenated (95% O2/5% CO2) for ≥30 minutes prior to brain slice preparation. Animals were anesthetized using isoflurane, after which the brain was perfused by clamping the aorta and injecting the slicing medium into the left atrium of the heart after which the right atrium was rapidly cut. Perfusion was maintained for approximately one minute, after which the brain was quickly removed. The brain was embedded in 2% agarose and coronal sections (300 *µ*m) were made using a VF-300 compresstome (Precisionary Instruments LLC). Slices were immediately placed in carbogenated artificial cerebrospinal fluid (ACSF; NaCl 120 mM; KCl 3.5 mM; MgSO4 1.3 mM; CaCl2 2.5 mM; D-glucose 10 mM; NaHCO3 25 mM; NaHPO4 1.25 mM) which was kept at 30°C. Upon placing the slices in ACSF, heating of ACSF was stopped and its temperature was allowed to drop to room temperature gradually for one hour before intracellular recordings where it was kept. All chemicals were obtained from Sigma-Aldrich unless otherwise specified.

Slice recordings were performed at room temperature in carbogenated ACSF (flow rate 1 ml/min) under a Nikon Eclipse FN1 upright microscope. Glass capillaries (Sutter GC150-15F; ID 0.5 mm; OD 1.0 mm) were pulled using a PP2000 pipette puller (Sutter) to prepare recording pipettes with an impedance of 8±2MΩ. During all experiments, pipettes contained the same intracellular solution (K-Gluconate 130 mM; KCl 5 mM; HEPES 10 mM; MgCl2 2.5 mM; Mg-ATP 4 mM; Na-GTP 0.4 mM; Na-phosphocreatine 10 mM; EGTA 0.6 mM). Fluorescence (mCherry) guided targeting was performed using a band pass filter (590 nm; Nikon G-2A) and a white LED light source (CoolLED). Data was acquired using a HEKA EPC9 amplifier controlled using HEKA PatchMaster software. Upon entering the whole-cell configuration, cells were kept stable at ∼-70 mV. The permanent magnet (cylindrical N42 Neodymium magnet, diameter: 1.6 mm) was positioned using a precision micromanipulator (Sensapex) within ∼200±100 *µ*m. The magnetic stimulation intensity at the recording site was calculated to be >140 mT (see Supplemental Figure 1 for magnetic intensity measurements).

Current clamp or voltage clamp protocols were performed with and without magnetic stimulation in every cell; the order was pseudo-random. Excitatory and inhibitory postsynaptic potentials were recorded for 2-5 minutes upon clamping each neuron 10 mV below its spiking threshold, which was determined using a step-and-hold protocol (t=500 ms, inter sweep interval=5 sec). If needed, current injection was adjusted to maintain the target membrane potential. In voltage clamp recordings, membrane potential was clamped 10 mV below the spiking threshold. To determine the membrane potentials at which voltage-gated conductances are initiated a triangular (sawtooth) stimulus protocol was used. The cell was clamped 10 mV below spiking threshold (V_start_) for 50 ms after which membrane potential was ramped linearly to -V_start_ over 100 ms before the membrane was depolarized back to V_start_, again over a 100 ms period. Each triangular pulse was repeated an additional 4 times after in every given sweep. Each sweep was repeated 3 times with an inter-sweep interval of 10 sec. Spontaneous excitatory and inhibitory postsynaptic currents were recorded while clamping neurons 10 mV below their spiking threshold for a duration of 2 – 5 minutes.

### Immunohistochemistry

After *in vivo* or *ex vivo* experiments, brain tissue (whole brains or acutely prepared slices) was fixed in 4% PFA at least overnight. The tissue was then transferred to a solution of 40% sucrose in phosphate buffered saline (PBS) until saturation, after which it was frozen using dry ice and cut to 40 µm sections using a Microm HM-430 sliding microtome (Thermo Scientific). Sections were individually stored in antifreeze at -30 °C until further use or transferred to PBS and immediately processed. For immunolabeling, sections were treated with the following solutions: 3× 20 minutes PBS; 1x 30 minutes 0.5% Triton-PBS; 2× 15 minutes PBS. Sections were then pre-incubated in blocking solution (PBS + 2% Normal Donkey Serum (Jackson $017-000-001) + 0.5% TSA Blocking Reagent (Perkin Elmer #FP1020) for 1 hour. Primary antibody (Rabbit anti-mCherry, Abcam #167453) was diluted 1:200 in blocking solution, and sections were incubated overnight. After primary antibody incubation, sections were washed 3x 15 minutes in PBS, followed by secondary antibody incubation. Alexa 488 Donkey anti-Rabbit (Jackson #711-545-152) was diluted 1:200 in blocking solution and sections were incubated for 3 hours. Secondary incubation was followed by 2x 15 minutes washing in PBS after which the sections were mounted onto glass slides and allowed to air-dry for 2 hours. Finally, FluorSave (Millipore #345789) was applied and sections were covered with a cover slip, and allowed to harden for at least 24 hours before confocal imaging. The imaging was done using the Leica SP8 inverted scanning confocal microscope using the LAS X software at the General Instrumentation Department of the Faculty of Science, Radboud University. Image processing was performed using Fiji.

Tetrode locations were confirmed using Nissl staining. After sectioning (coronal; 40 *µ*m), sections were mounted onto gelatin-coated slides and left to air dry. Slides were then transferred to 96% ethanol for 10 minutes, followed by sequential steps (2 min/each) of 90%, 80%, 70%, 50% ethanol and finally rinsed in demineralized water. Tissue was then stained using 0.1% cresyl fast violet in demineralized water for 30 minutes, followed by 5 minutes washing in demineralized water. Sections were then dehydrated by rinsing in sequential steps of 50%, 70%, 80%, 90% and 96% ethanol (2 min/each). Differentiation was done in acidified 100% ethanol for 1 minute. For mounting, slides were transferred to 100% ethanol for 2 minutes, followed by 2× 5 minutes xylene after which mounting medium (Entellan) was applied and slides were covered with a glass coverslip.

### Culturing and transfection of Neuro-2a (N2a) cells

Neuro2a (N2a) cells were obtained from ATCC (CCL-131) and maintained in culture medium (MEM (Gibco #41090-028) supplemented with 10% FCS (Gibco #10270), Na-Pyruvate (Gibco #11360-039) and Pen/strep (Gibco #15140-122)) at 37 °C and 5.5 % CO_2_ atmosphere. One day prior to transfection, 25,000 cells were seeded onto 14-mm coverslips in 24-wells plates (for immunofluorescence assay) or 50,000 cells per well in a 12-wells plate (for Western blot analysis). For transfection jetPRIME (Polyplus Transfection #114-15) transfection reagent and endotoxin-free plasmid DNA (Macherey-Nagel Nucleobond #740422.10) were used. Recombinant protein expression was expressed for 48 hours before biochemical and immunohistochemical assays were performed.

### Immunofluorescence assay for N2a cells

Cells were washed with PBS, fixed for 30 min with 4% paraformaldehyde in PBS at 4 °C, and re-washed with 50 mM NH_4_Cl/PBS to block any remaining PFA residuals. Cells were permeabilized using PBS-T (PBS/0.1% Triton-X100), blocked for 30 min in blocking buffer (PBS-T/2% BSA) at RT and incubated o/n at 4°C with mouse anti-Flag tag antibody (ThermoFisher #MA1-91878, 1:100 in blocking buffer). After washing with PBS-T, cells were incubated with secondary antibody Goat-anti-mouse IgG (H&L)-Alexa488 (Invitrogen #A11001, 1:500 in blocking buffer) for 1 hour at RT, washed with PBS-T, PBS and incubated in 300 nM DAPI (Invitrogen #D-1306) in PBS for 5 min to stain the nuclei. Following a final washing step with PBS, cells were embedded using FluorSave (Calbiochem # 345789 #). Imaging was performed using a Leica SP8x confocal laser scanning microscope.

### Western blot analysis

Cells were lysed in 200 *µ*l lysis buffer (150 mM NaCl, 1.0% Triton-X100, 0.5% sodium deoxycholate, 0.1% SDS, 50 mM Tris, pH8.0 supplemented with protein inhibitor mix (cOmplete Roche # 11873580001) and incubated on ice for 10 min. The lysates were cleared by centrifugation (12,000 g, 10 min at 4 °C) and 20 *µ*l of cell lysate was denatured using Laemmli sample buffer for 4 min at 100 °C, separated on a 10% SDS-PAGE and subsequently transferred onto PVDF membrane (Hybond, Amersham). After blocking, the membranes were incubated with anti-flag tag (1:500), anti-mCherry (1:1000) and anti-tubulin (E7, 1:100) primary antibodies o/n at 4 °C. Secondary HRP-conjugated Goat-anti-Rabbit (Santa Cruz # sc-2004, 1:5000) or Goat-anti-Mouse (Santa Cruz # sc-2005, 1:5000) antibodies were used, followed by chemiluminescence (Pierce, ECL plus). Signal detection was done using an ImageQuant LAS 4000 (GE Healthcare) imaging system.

### Data analysis

All data analyses were performed offline in Matlab using custom-written software, unless stated otherwise.

### In vivo recordings

To facilitate spike detection, the extracellular signal was zero-phase bandpass filtered between 600 and 6000 Hz. Outliers in the frequency domain were removed via an adaptive filter (Jun et al., 2017) before spike detection and sorting were performed using KiloSort (Pachitariu et al., 2016). If necessary, the resulting clusters were merged. The normalized distance in principal component space between any two clusters projected on the axis joining their two centroids was used as a merging criterion as described (Yger et al., 2018). To obtain more accurate merging results, high frequency oscillatory noise that is sometimes present, due to e.g. electrical interference, is substituted with Gaussian noise based on the background signal. Additionally, spike clusters with a mean spike amplitude of less than 4 times the standard deviation of the background noise are removed before doing principal component analysis and cluster merging. Note that unlike other clustering algorithms KiloSort does not include the thresholding pre-processing step, thus spikes with smaller amplitudes are also detected (albeit they tend to be multi-unit clusters).

The standard deviation of the background noise, σ, was estimated from the mode of the signal envelope, given by the magnitude of the analytical signal calculated through Hilbert transform (Dolan et al., 2009). High amplitude stimulus artifacts contaminating the spike clusters were deleted from the cluster depending on the mean-squared error between the waveform of the potential artifact and the mean waveform of the cluster.

To assess the quality of each cluster, a number of quality metrics were computed (see Supplemental Figure 2). Clusters were automatically classified based on these metrics as either being “noise”, a “multi-unit”, a “contaminated single unit” or an “isolated single unit”. Only isolated single units are considered for the downstream analysis. In order to be labelled as an isolated single unit, the cluster must pass the criteria listed in Supplemental Table 2. These criteria include common spike metrics as well as numerical description of the temporal stability of the cluster and waveforms correlations across channels. To determine the temporal stability, it is assumed that the neuron’s firing pattern can be *roughly* approximated by a homogeneous Poisson process. Strong deviation from this process is typically an indication that the cluster is highly contaminated with mechanical or stimulus artifacts that only occur at specific moments during the session, leading to a (very) irregular firing pattern. This instability is quantified by counting the number of spikes in 10 second intervals over the entire session and fitting the resulting distribution with a Gaussian (Supplemental Figure 2d). The integral of the distribution predicted by the Gaussian is then computed as the measure for the temporal stability. A Gaussian with standard deviation λ^3/2^ (where λ is the event rate) was used for the fit. This was empirically determined as it better deals with typical inhomogeneities in the Poisson process at higher firing rates compared to a regular Poisson distribution. Lastly, to determine the single unit isolation quality, a mixture of drifting t-distributions is fitted to the spiking data (Shan et al., 2017), which returns the fraction of false positive and negative spikes in each cluster. A toolbox has been developed in Matlab that carries out the aforementioned steps to process the extracellular data (Brouns and Celikel, 2019). The source code can be downloaded from https://github.com/DepartmentofNeurophysiology/Paser.

FieldTrip (Oostenveld et al., 2011) was used to construct interspike interval histograms (ISIH), (joint) post-stimulus time histograms (jPSTH) and correlograms, which were compared between magnet-on and magnet-off (control) conditions. The jPSTHs were normalized by subtracting the product of the individual firing rates from the joint firing rate and divided by the product of the two standard deviations (Aertsen et al., 1989). A bin size of 0.5 ms was selected for the ISIHs and 1 sec for the (j)PSTHs.

To separate the low frequency local field potential (LFP) activity from the high frequency (spiking) components, the raw signal is low-pass filtered (<300 Hz) before a second order IIR notch filter was applied to eliminate the 50 Hz mains hum. The resulting data was downsampled to 1.2kHz. Artifacts due to mechanical motion of the drive and electromagnetic interference were removed by locating uncharacteristically large peaks in the filtered signal amplitude, its derivative and power spectral density (Bakštein et al., 2017) using twice the median absolute deviation as an automatic threshold, and then replacing the detected artifacts with a linear interpolation between neighbouring sample points. Time-frequency analysis was subsequently carried out using FieldTrip (Oostenveld et al., 2011). Power-spectra were estimated by fast Fourier transform after Hann tapering the signal, followed by spectral smoothing with a Gaussian kernel.

### In vitro recordings

Intracellular recordings included both current-clamp and voltage-clamp recordings. In current-clamp configuration a step-and-hold protocol was used to determine the pattern and statistics (e.g. rate, timing, spike threshold) of action potentials. Excitatory (EPSP) and inhibitory post-synaptic potentials (IPSP) were recorded separately in the absence of any somatic current injection. In the voltage-clamp configuration triangular pulse injections were used to determine at which membrane voltage the conductances are observed. Excitatory (EPSC) and inhibitory (IPSC) post-synaptic currents were recorded in the absence of any voltage-clamp without tetrodotoxin in the bath. All recordings included within cell controls, i.e comparisons were across magnet-on vs magnet-off conditions. The data was split into groups post-hoc before they were treated as described below.

For detection of EPSC and IPSC events, the signal was denoised using a second order IIR notch filter at 50Hz. To detect the inward currents, the filtered signal was first smoothed by convolving with a Gaussian kernel, then binned (1 sec/bin). The mean and standard deviation of the processed signal in every interval were computed separately, and used to set a threshold for detection of candidate events. A second threshold, calculated after excluding the candidate events determined upon first thresholding, allowed adapting the event detection threshold and ensured that the event threshold estimate is not confounded by the rate of activity in any given period. The candidate events identified after this second passage were classified as events if they satisfied selection criteria including: 1) duration of the events is 2 < x < 1000 ms, 2) the absolute peak appears within the ⅓ of the event duration, 3) the peak duration is <1.5 ms, 4) the current decays with an exponential decay from the absolute peak, 5) the current decay is complete by the end of the event as quantified by comparing the average current within the first 0.5 ms of the event to the last 0.5 ms. Events with a decay rate < 0 (ms^-1^), and integral < 5 (pA*ms) were excluded from the analysis. The characteristics of each event: amplitude, duration, decay rate (estimated with fitting an exponential decay), total current, instantaneous frequency, were then computed using the raw, non-smoothed signal. The signal processing was identical for the inward and outward currents. In the case of inward current, the data was inverted before event detection. For detection of EPSP and IPSP events the signal was denoised using a second order IIR to remove the 50Hz mains hum, detrended and a polynomial was fit to the data. Events were detected using the “findpeaks” function in the Signal Processing Toolbox of Matlab with minimum peak amplitude of 1 mV. Interpeak interval was > 2ms. The signal processing was identical for the EPSP and IPSP events. In the case of IPSPs, the data was inverted before event detection. For stimulus evoked response detection, as for ramp-and-hold stimulation protocol in voltage-clamp and current-clamp configurations, and the triangular sweeps in voltage clamp, evoked responses were detected using “findpeaks”. The minimum peak amplitude was defined as 30 mV in current clamp and 0.2 nA in voltage-clamp configuration during the ramp-and-hold protocol. Otherwise it was set to 0.1 nA. In voltage-clamp configurations event detection was performed in the absolute signal.

### Statistical analysis

Paired t-test was used in comparisons between within cell across stimulation comparisons, unless otherwise stated. When there are more than one independent variable, two-way ANOVA was used after testing for normality and homoscedasticity of the distributions.

## Acknowledgements

We thank the members of the Department of Neurophysiology for stimulating discussions and the critical insight on the manuscript. This work was supported by grants from the European Commission (Horizon2020, nr. 660328), European Regional Development Fund (MIND, nr. 122035) and the Netherlands Organisation for Scientific Research (NWO-ALW Open Competition, nr. 824.14.022) to TC, as well as doctoral fellowships from the Chinese Scholarship Council to YZ and XY, and the National Council for Scientific and Technological Development of Brazil (CNPQ) to ASL.

## Supplemental Figures

**Supplemental Figure 1.**
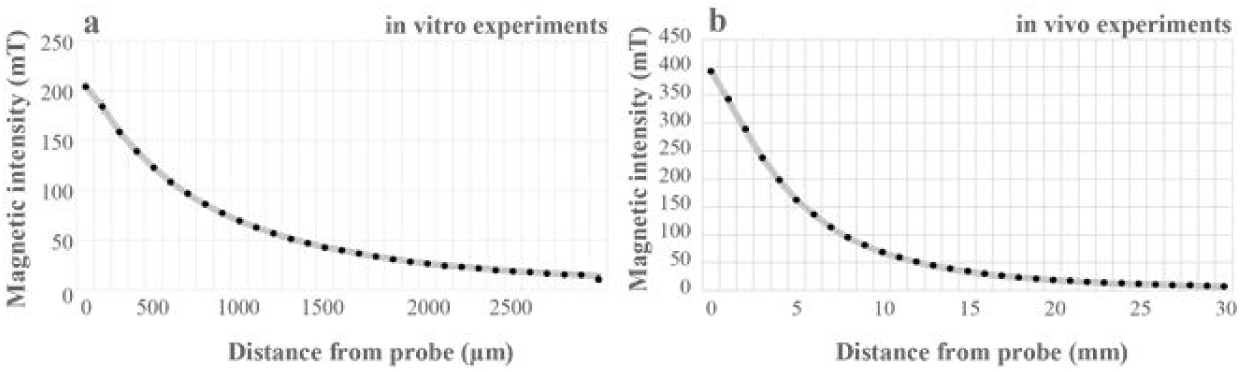
Change in magnetic field strength as a function of distance from the permanent magnet. Measurements were made using a HT-20 Gaussmeter (Hangzhou BST Magnet Co. Ltd., China) and repeated three times at each distance. The error bars are standard deviation. The distance between the target cells and magnet was ∼200±100 *µ*m in in vitro and ∼8±2 mm in in vivo experiments.

**Supplemental Figure 2.**
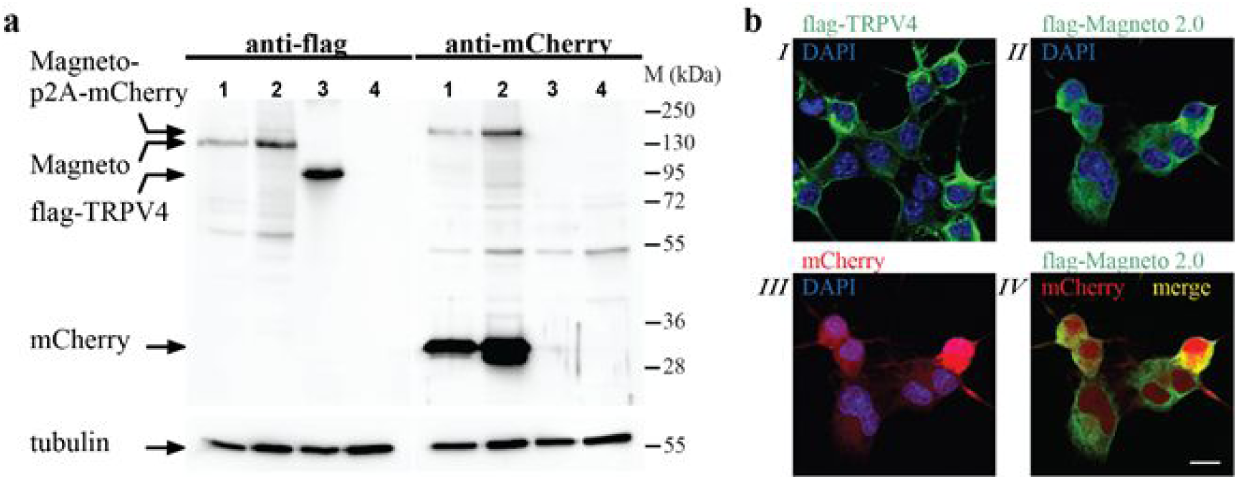
Magneto2.0 is cleaved from mCherry but remains primarily in the cytoplasm. (a) To confirm that Magneto was effectively released from its fusion with mCherry, N2A cells were transfected with 1: pFUGW-V4trunc-fer-traffick-p2A-mCherry, 2: pcDNA3-Magneto-p2A-mCherry, 3: pcDNA3-flag-TRPV4 and 4: pcDNA3 plasmids. TRPV4 and empty pcDNA3 served as controls; tubulin was used as a loading control. Magneto and TRPV4-expression were analyzed by Western blot using anti-flag and anti-mCherry antibodies. The major protein products identified by the anti-flag antibody were the ∼135 kDa Magneto protein and the ∼95-kDa flag-TRPV4 protein. The ∼160-kDa Magneto-p2A-mCherry product was recognized by both anti-flag and anti-mCherry antibodies. However, the principal protein detected by the anti-mCherry antibody was mCherry. These findings indicate that Magneto was effectively cleaved from its fusion with mCherry. Note that both pFUGW-V4trunc-fer-traffick-p2A-cherry and pcDNA3-Magneto-p2A-mCherry produce the same protein products, indicating that sub-cloning of Magneto into the viral vector pFUGW did not affect the Open Reading Frame of Magneto-p2A-mCherry. (b) I: N2a cells were transfected with pcDNA3-flag-TRPV4 and immunostained using anti-flag antibodies. Note that flag-TRPV4 was observed in the plasma membrane. II-IV: N2a cells were transfected with Magneto-p2A-mCherry plasmid and stained using anti-flag antibodies (II, IV, green). mCherry fluorescence was directly imaged (II, red), DAPI (blue) was used to stain the nuclei (II-III). Note that mCherry was mostly localized to the nucleus, however, also a significant amount was present in the cytoplasm. In contrast to flag-TRPV4, most Magneto expression was found in reticular structures in the cytoplasm, most likely representing the ER (IV). These results indicate that, compared to flag-TRPV4, Magneto is less effectively transported to the plasma membrane of N2a cells. Scale bar: 10 µm.

**Supplemental Figure 3.**
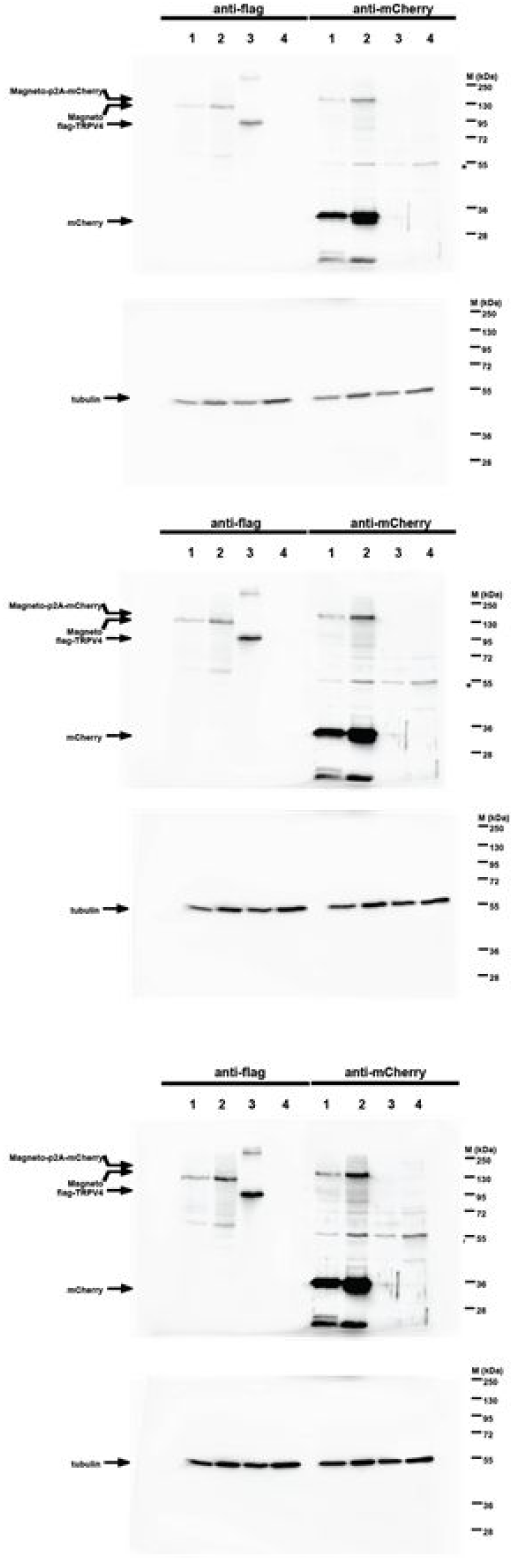
The Western Blot shown in Supplemental Figure 2a at different exposures.

**Supplemental Figure 4.**
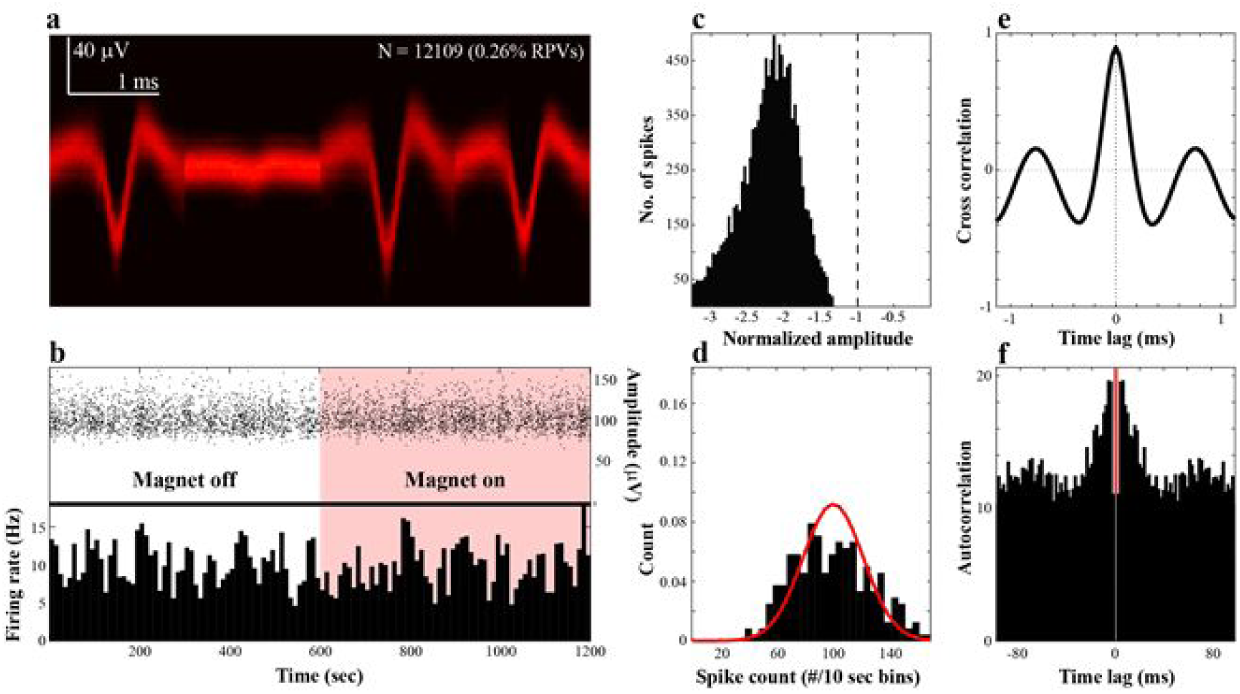
Isolation quality of single units. (a) Concatenated spike shapes across the tetrode channels visualized as a density heatmap of time-voltage values. (b) The peak-to-peak amplitude (top) and firing rate (bottom) over the whole recording period in a session, showing the temporal stability of the cluster. (c) Histogram of normalized spike amplitudes. Any amplitude to the left of the dashed vertical line crosses the minimum threshold for spike detection, which is three times the average background signal (voltage). (d) Distribution of running average spike counts in 10 sec bins (with 5 sec overlap). The red line indicates the predicted distribution of spike counts by a Gaussian distribution given the observed variance in spike count. (e) Cross-correlation between two tetrode channels that are maximally dissimilar. The large peak at zero lag indicates that the spike waveforms are temporally aligned. (f) Autocorrelation of spike events.

**Supplemental Figure 5.**
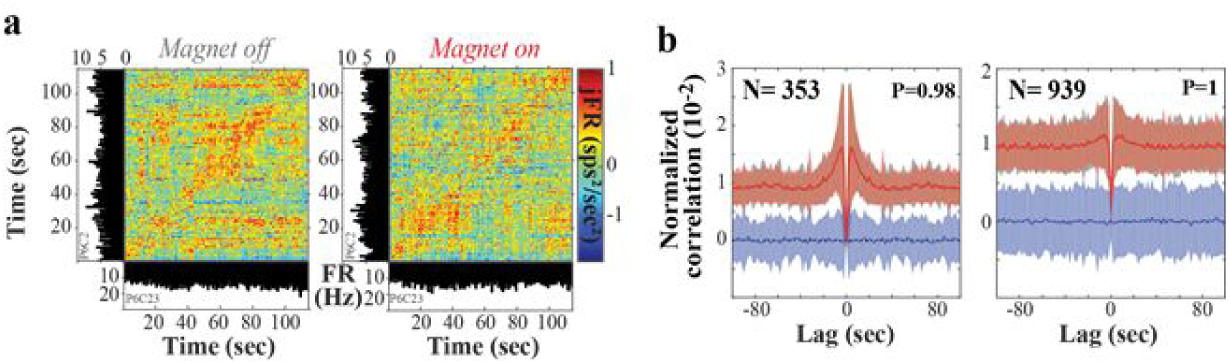
Magnetic stimulation does not alter the temporal correlations of action potentials. (a) A representative example of spiking correlations between two simultaneously recorded neurons from the same tetrode. Joint peristimulus time histograms represent the temporal correlations across the binary states of magnetic stimulation. (b) Spiking pattern in single units (auto-correlations; left) and across simultaneously recorded units (cross-correlations; right). Grey traces: Correlation in the absence of magnetic stimulation (magnet off), red: during magnetic stimulation (magnet on), blue: the pairwise difference between magnet off-magnet on. Thick traces are population averages; color-coded shadows in the background represent the standard deviation within stimulation condition. Neither single cell spiking correlations (N=353, P=0.98, paired t-test), nor spiking correlations across neurons (N=939, P=1.00, paired t-test) were altered upon magnetic stimulation. See Supplemental Figure 5 for Poincaré analysis of spiking in single neurons across stimulus conditions.

**Supplemental Figure 6.**
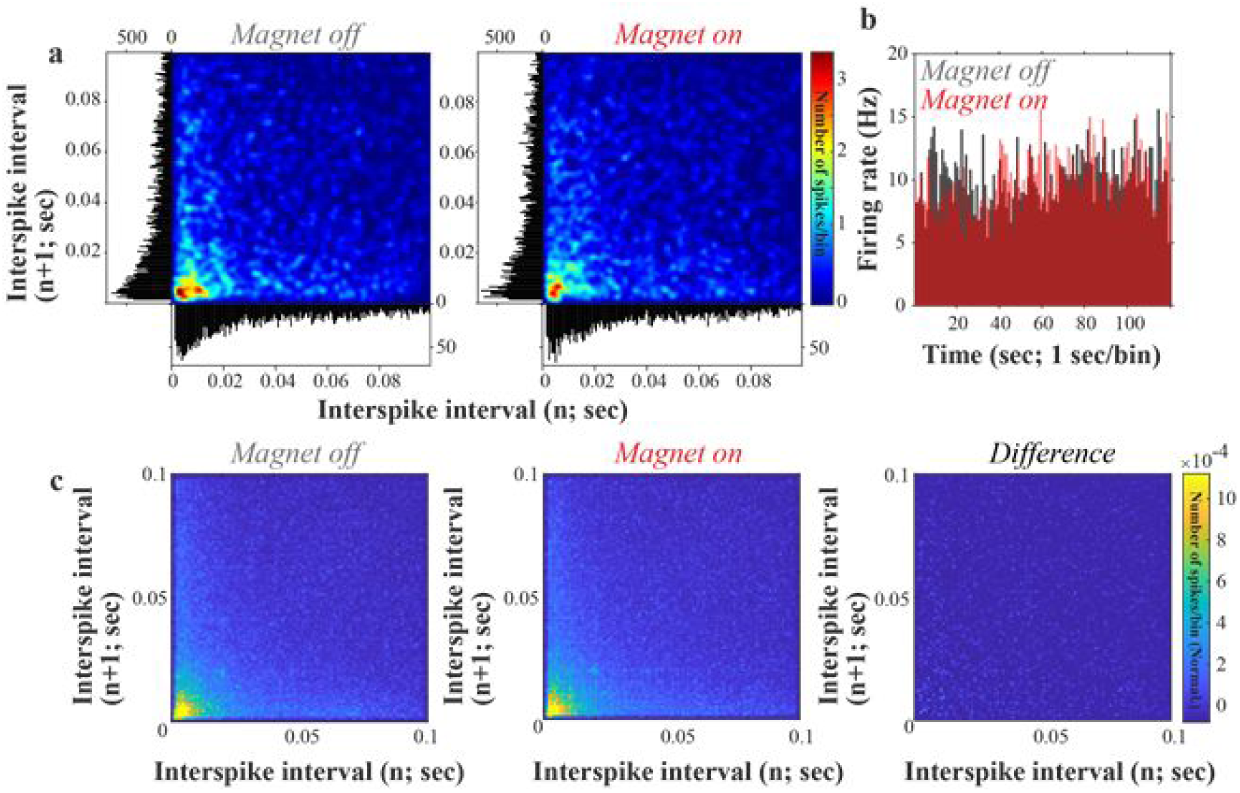
Spiking pattern does not change during magnetic stimulation in vivo. (a) Poincaré plot of interspike intervals (ISI) from a representative neuron under control conditions (magnet off) and during magnetic stimulation (magnet on). Corresponding ISI histograms (bin size = 0.5 ms) are shown on the left and bottom of each plot. (b) Post-stimulus time histograms of a representative neuron for the magnet on and off conditions. (c) Mean normalized ISI Poincaré plot across all neurons (N = 235). Neurons that fired <50 spikes during the period of observation were excluded from the plot (N = 118).

**Supplemental Figure 7.**
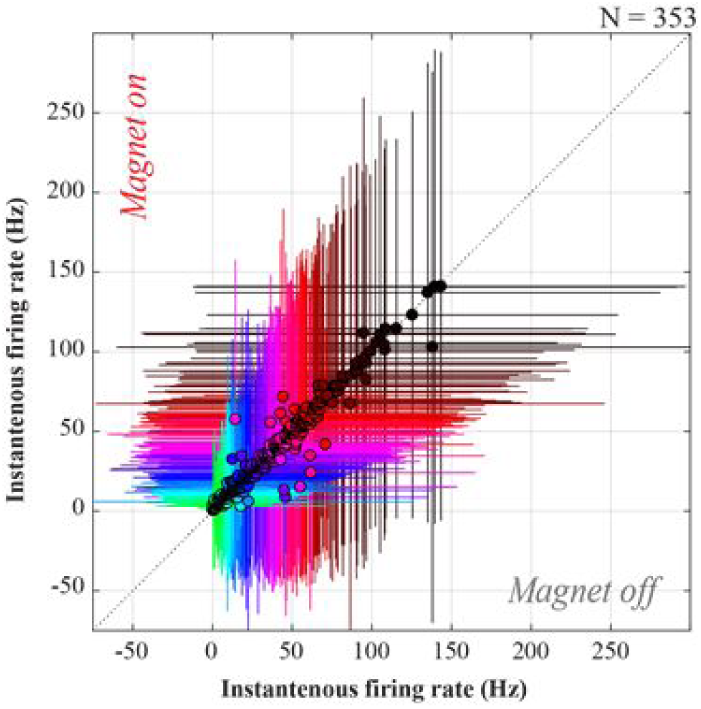
Firing rate of identified single neurons before and during magnetic stimulation. Each dot represents the average firing rate of a neuron (N=353) across the two conditions as in Figure 1c. The color code represents firing rate. The error bars are standard deviation from the mean within session. See Figure 1c for the results of statistical comparison.

---

**Supplemental Table 1.**
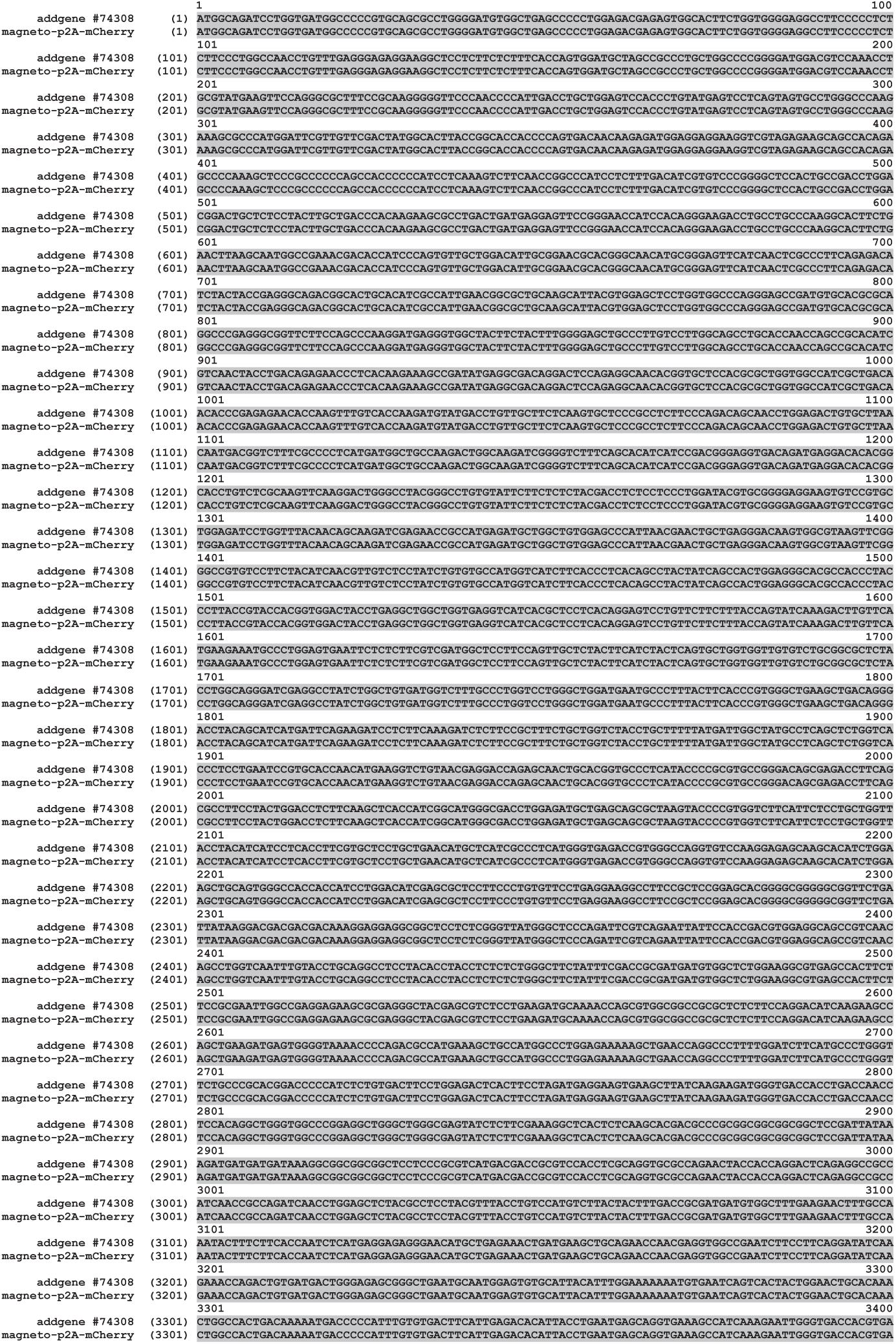
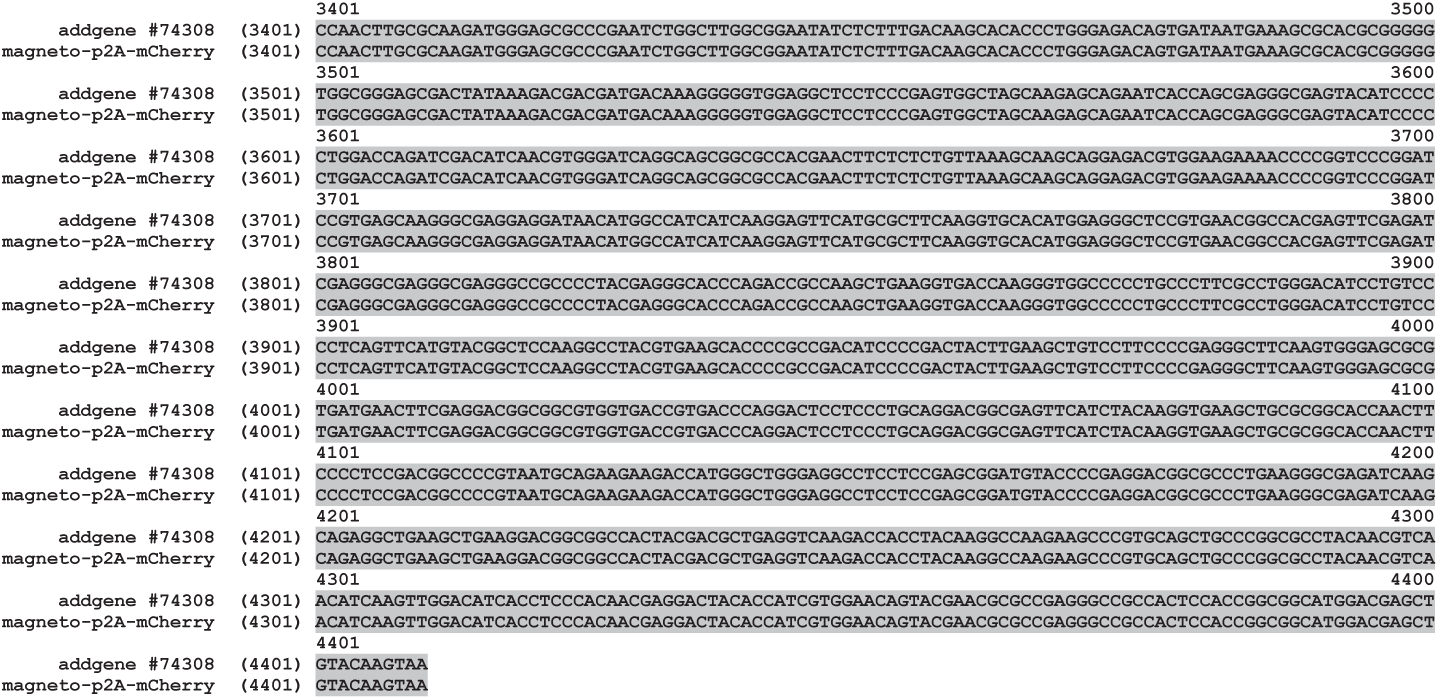
Sequence comparison between the original plasmid pcDNA3.0-Magneto2.0-p2A-mCherry (Addgene, #74308) and the open reading frame of the lentiviral vector used for viral gene transfer in this study.

**Supplemental Table 2.** Quality criteria to determine whether a cluster is a single unit and well isolated. Level of background noise is indicated by σ.

| Single unit quality metrics | Threshold |
| --- | --- |
| Number of spikes | > 50 |
| Mean relative spike amplitude | > $4\sigma$ |
| Mean absolute spike amplitude | < 300 μV |
| Percentage of spikes with an amplitude less than $3\sigma$ | < 10% |
| Percentage of refractory period violations (< 1.5 ms inter-spike interval) | < 5% |
| Temporal stability measure of spike counts in 10 sec interval | > 0.4 |
| Location of cross-correlation peak between channel waveforms | < 0.20 ms |
| **Isolation quality metrics (mixture of drifting t-distributions)** |  |
| Percentage of false-positive events | < 10% |
| Percentage of false-negative events | < 10% |

